# The neural mechanisms of active removal from working memory

**DOI:** 10.1101/2021.03.16.435617

**Authors:** Jiangang Shan, Bradley R. Postle

## Abstract

The ability to frequently update the contents working memory (WM) is vital for the flexible control of behavior. Whether there even exists a mechanism for the active removal of information from working memory, however, remains poorly understood. In this Registered Report we will test the predictions of models for two different (and not mutually exclusive) mechanisms of active removal: adaptation-hijacking and mental-context shifting. We will collect functional magnetic resonance imaging (fMRI) data while subjects perform a novel “ABC-retrocuing” task designed to elicit two modes of removal, active or passive (Shan & Postle, Registered Report). The adaptation-hijacking model posits an adaptation-like modification of perceptual circuits combined with a weak activation of the to-be-removed item. Its predictions will be assessed by using multivariate inverted encoding modeling (IEM) and photic “pings” to assay the state of feature-selective encoding channels and of activity-silent representations under active-removal versus passive-removal conditions. A second model – “working memory episodic memory” (WMEM) – posits that interference from no-longer-relevant information is minimized by making the mental context associated with new information dissimilar from that associated with the to-be-“removed” information. This will be tested by using representational similarity analysis (RSA) to compare the rate of contextual drift under active-removal versus passive-removal conditions.

## Introduction

A hallmark of working memory (WM) is that it is rapidly updateable, such that information that was relevant in the recent past can be easily replaced once circumstances change and different information is now of primary importance. One way that this is operationalized in the laboratory is with a block of stand-alone trials (e.g., of delayed recognition): Once trial *n* has been completed, subjects have little difficulty encoding a new memory set for trial *n + 1*. Because the set of items is randomly selected for each trial, the memory items for each trial lose their relevance at the end of that trial, and our intuition tells us that they should be removed from WM. However, the phenomenon of proactive interference indicates that the assumed removal of no-longer-relevant information is often not complete. This is particularly notable for trials featuring “recent-negative” recognition probes that were not in the memory set of the current trial, but were in the memory set on the previous trial; these lead to an increased false-alarm rate, and longer reaction times (RTs) for correct rejections (Monsell, 1978). For visual WM tasks that test recall, the imperfect nature of removal manifests itself as serial dependence. For example, when the orientation of a Gabor patch is the feature to memorize and recall, the reported orientation for trial *n* is commonly found to be attracted toward the orientation of the item that had been shown on trial *n - 1* (e.g., Bliss et al., 2017; Fischer & Whitney, 2014).

There is also a growing body of neural evidence for the incomplete removal of information from WM. In an electroencephalography (EEG) study of delayed recall of orientation, Bae and Luck (2019) were able to decode the orientation of the previous trial’s sample after the onset of the current trial’s sample. For delayed recall of location, Barbosa, Stein et al. (2020) were able to decode the previous trial’s sample location (from activity in the PFC of nonhuman primates) from late in the intertrial interval (ITI), just prior to the start of the next trial. A similar pattern of reactivation was also observed in whole-scalp EEG in humans. Simulations with a bump-attractor network model suggested that the reactivation of no-longer relevant information may be due to “nonspecific” activation of a residual neural trace that is “imprinted in neuronal synapses as a latent activity-silent trace” (Barbosa, Stein, et al., 2020). Tellingly, this model did not include an explicit mechanism for removal of no-longer-relevant information; rather, when an item was no longer relevant, activation was simply withdrawn from it, and the bump of activity representing it receded to baseline. Similarly, another formal model of WM performance, the Prefrontal Basal Ganglia Working Memory (PBWM) model, accomplishes the replacement of a no-longer-relevant item with a new one via the “reallocation” of resources away from the former (Chatham and Badre, 2013). Thus, there is considerable evidence that a default strategy for updating the contents of WM is what we will refer to as the “passive removal” of no-longer-relevant information.

In contrast to passive removal, there is also considerable evidence for an active removal mechanism, particularly during tasks that require the simultaneous maintenance of multiple items in WM. One example is dual serial retrocuing (DSR) tasks, in which subjects are first shown two stimuli to memorize, then a retrocue indicates which will be tested first; after the first test, a second retrocue is shown to indicate (with equal probability) which of the two items will be tested in a second test. The first retrocue designates one of the two as a “prioritized memory item” (PMI), and the uncued item, by default, becomes an “unprioritized memory item” (UMI). Critically, the UMI can’t be removed from WM, because it may be needed for the second test. After the second retrocue, the cued item takes on the status of PMI, and the uncued becomes irrelevant for the remainder of the task (an “irrelevant memory item,” IMI). In functional magnetic resonance imaging (fMRI) and EEG studies, multivariate evidence for an active trace of the UMI often drops to baseline (e.g., LaRocque et al., 2013; Lewis-Peacock & Postle, 2012), but the fact that it has not been removed from WM is inferred from that fact that a pulse of transcranial magnetic stimulation (TMS) produces a transient reactivation of its active trace (Rose et al., 2016), as well as an increase in false alarm responding when the UMI is used as an invalid memory probe (reminiscent of proactive interference effects; Rose et al., 2016; Fulvio & Postle, 2020). Evidence for the active removal of the IMI, in contrast, is inferred from the absence evidence for either TMS reactivation (Rose et al., 2016) or a TMS-related false alarm effect (Rose et al., 2016; Fulvio & Postle, 2020).

What might be the mechanism of the active removal of information from WM? In the interference model of visual WM (Oberauer & Lin, 2017), active removal is accomplished by breaking the association between the content of the memory item (e.g., the orientation of a Gabor patch) and its context (e.g., where this patch had been presented on the screen; Lewis-Peacock et al., 2018). In a bump attractor model, such as that of Barbosa, Stein et al. (2020), the active removal of a memory item could be effected with a strong nonspecific burst of activity that would saturate the network, effectively erasing any trace of the IMI. A third possibility comes from a recently articulated “Working Memory Episodic Memory” (WMEM) model, which proposes that “flexible forgetting” from WM is accomplished by shifting the mental context with which more recent information is encoded, thereby making the retrieval of the IMI less likely (Beukers et al., 2021). Although the design of the present Registered Report will allow us to assess evidence for two of these hypothesized mechanisms, its primary motivation is to evaluate a novel mechanism that we will introduce next: the adaptation-like modification of perceptual circuits.

### Behavioral evidence for the active removal of information from WM

In a recent behavioral study, Shan and Postle (Registered Report) designed a novel “ABC-retrocuing” task intended to engage active or passive removal of an IMI from WM. Each trial began with the simultaneous presentation of two sample oriented gratings (items *“A”* and *“B”*) in two of six possible locations. After a brief delay, a circle appearing at one of the two locations indicated that the corresponding item (for this example we’ll say *A*) might be tested at the end of the trial, thereby designating *A* a PMI and *B* an IMI. After another brief delay, a third item (“*C*”) was presented, and at the end of the trial recall of the orientation of either *A* or *C* was tested with a response dial appearing at the location of the to-be-recalled item. The critical manipulation that was intended to encourage active versus passive removal was the location at which item *C* would be presented: In the *overlap* condition, item *C*’s location was always the same as that of the IMI (i.e., item *B*); and in the *no-overlap* condition item *C* always appeared at one of the locations that had not been occupied by either item *A* or *B*. Trials were blocked by condition, and subjects were explicitly informed about the condition prior to each block. The logic was that the *no-overlap* condition might encourage passive removal, just because this seems to be the default for many working memory tasks, as evidenced by proactive interference and serial dependence effects. For the *overlap* condition, however, subjects might be motivated to actively remove the IMI from WM, because otherwise its shared location with item *C* would lead to retrieval conflict when the response dial appeared at this shared location (i.e., “cue conflict”; Oberauer & Lin, 2017). A final element of the procedure is that each ABC-retrocuing trial was followed by a trial of simple 1-item delayed recall of orientation, with serial dependence of the latter on the immediately preceding ABC-retrocuing trial used to index the fate of the IMI.

Preliminary results from Shan and Postle (Registered Report) revealed a striking difference between the *overlap* and *no-overlap* conditions. In the *no-overlap* condition, item *B* had an attractive serial bias on 1-item recall, consistent with the attractive serial bias observed in several previous studies that we assume were characterized by passive removal of no-longer-relevant stimulus information (e.g., Bliss et al., 2017; Fischer & Whitney, 2014; Fritsche et al. 2020). In the *overlap* condition, in contrast, item *B* had a repulsive serial bias on 1-item recall, suggesting that it was processed in a very different way. The interpretation that the critical difference between the two conditions was active versus passive removal of the IMI was reinforced by the fact that in both conditions item *A* exerted comparable levels attractive serial bias on 1-item recall.

### Do common mechanisms underlie repulsive serial dependence and active removal from WM?

Recent work from three independent groups has converged on the view that serial bias effects arise from opposing effects that play out at two levels of processing: at the level of perception, adaptation to recent perceptual events produces repulsion from previous stimuli; at the level of decision making, in contrast, perceptual decisions are attracted toward previous decisions (Pascucci et al. 2019; Fritsche et al. 2020). Although decisional biases are stronger, explaining why short-lag serial bias is typically attractive, perceptual adaptation is longer lasting, explaining why longer-lag serial bias (e.g., the influence of the item from five trials previous) can be repulsive (Fritsche et al., 2020). That perceptual adaptation-like effects might also be involved in processing stimuli during WM comes from a simulation of fMRI results from a retrocuing task (Lorenc, Vandenbroucke et al., 2020). The observed effect of a retrocue on the multivariate inverted encoding model (IEM) reconstruction of the IMI was to “flip” it (c.f., Sahan et al., 2020), and this effect was best modeled by an adaptation-like suppression of the gain of perceptual feature channels corresponding to the value of the IMI, combined with an intermediate level of activation of the memory representation. Integrating across these results from the perceptual decision-making (Pascucci et al. 2019; Fritsche et al. 2020) and WM (Lorenc, Vandenbroucke et al., 2020, Shan & Postle, Registered Report) literatures has given rise to the idea that is the motivation for this Registered Report: The active removal of information from WM may be implemented via a top-down “hijacking” of an adaptation-like modification of perceptual circuits. More specifically, removal may be accomplished by the co-occurrence of two events. The first is the adaptation-like modulation of the gain of the perceptual channels that were engaged by the encoding of the IMI. This putative operation is “adaptation-like” because it is triggered by the onset of the retrocue (not by the perceptual processing of the to-be removed item) and because it is greater in magnitude than true adaptation (the repulsive effects of true adaptation are typically weaker than the attractive influence of recent decisions). The coincident second event is the brief, weak activation of the IMI. Although seemingly paradoxical, the weak activation of the to-be-removed item is predicted by the simulation from Lorenc et al. (2020) and is consistent with other computational accounts of forgetting (e.g., Kim et al., 2020).

We plan to assess this “hijacked adaptation” idea by collecting fMRI data while subjects perform an ABC-retrocuing task (Figure 1) while high-contrast task-irrelevant visual stimuli are flashed to “ping” the visual system, so as to assay predicted consequences of this hypothesized mechanism for active removal. Here we will provide a narrative overview of the logic of this experiment, to provide context for the statement of *Preregistered Hypotheses*. Details follow in the *Methods*.

**Figure 1.**
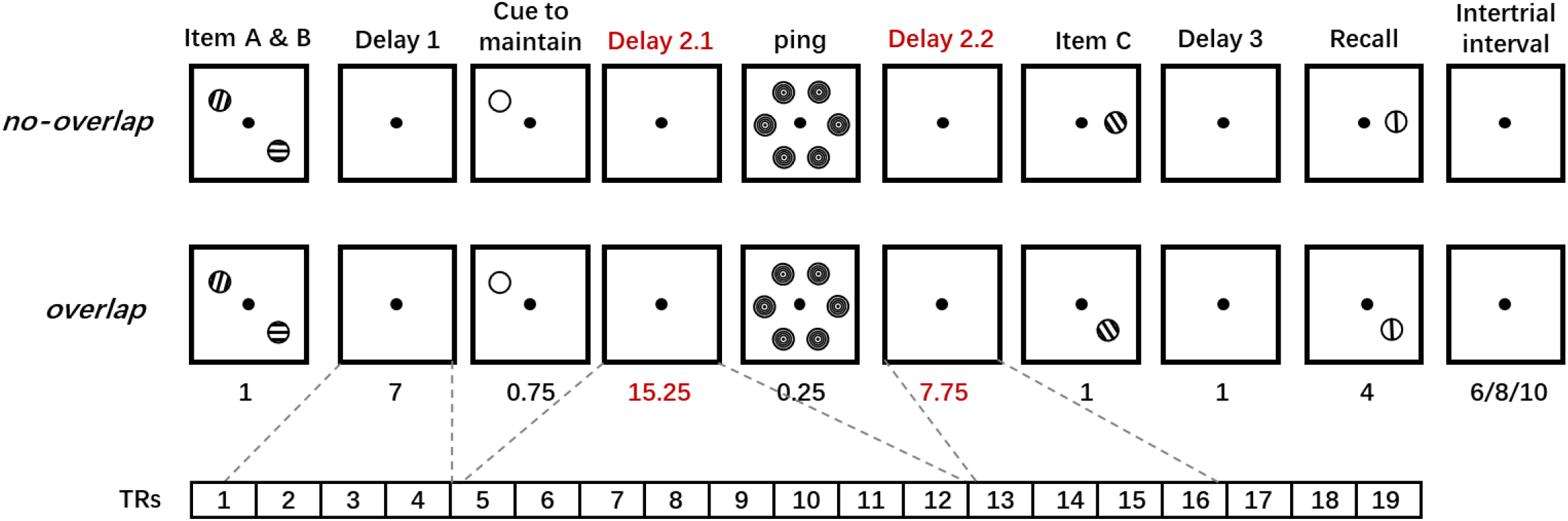
The experimental procedure.

Based on Shan & Postle (Registered Report), we assume that the IMI undergoes active removal in the *overlap* condition of the ABC-retrocuing task, but passive removal in the *no-overlap* condition. As diagrammed in Figure 2, in the *overlap* condition, we predict that the hypothesized adaptation-hijacking operation will produce a phasic “flipping” of the active representation of the IMI (operationalized as an IEM reconstruction of the IMI with a negative slope) during the first several seconds following the retrocue (c.f., Lorenc et al., 2020), followed by a disappearance of a detectable active trace (i.e., an IEM reconstruction slope not different from 0). This will correspond to successful removal of the IMI. A longer-lasting consequence of active removal, however, will be the residual adaptation-like change to the perceptual feature channels that correspond to the orientation of the IMI. This will be revealed in the filtering of the ping-evoked response, which will also produce a transient flipped IEM reconstruction of the IMI. Note that the delay period after the retrocue and before the ping will be relatively long (15.25s), so as to be able to dissociate the endogenously generated flipped reconstruction of the IMI that is triggered by the retrocue from the flipped reconstruction predicted to be evoked by the ping. In the *no-overlap* condition, we predict that the withdrawal of attention will result in the disappearance of evidence for an active representation of the IMI during the first several seconds following the retrocue. However, because the activity-silent trace of the IMI will not have been removed, the ping-evoked response will produce a conventional (i.e., not flipped) IEM reconstruction of the IMI (c.f., Barbosa, Stein, et al., 2020).

**Figure 2.**
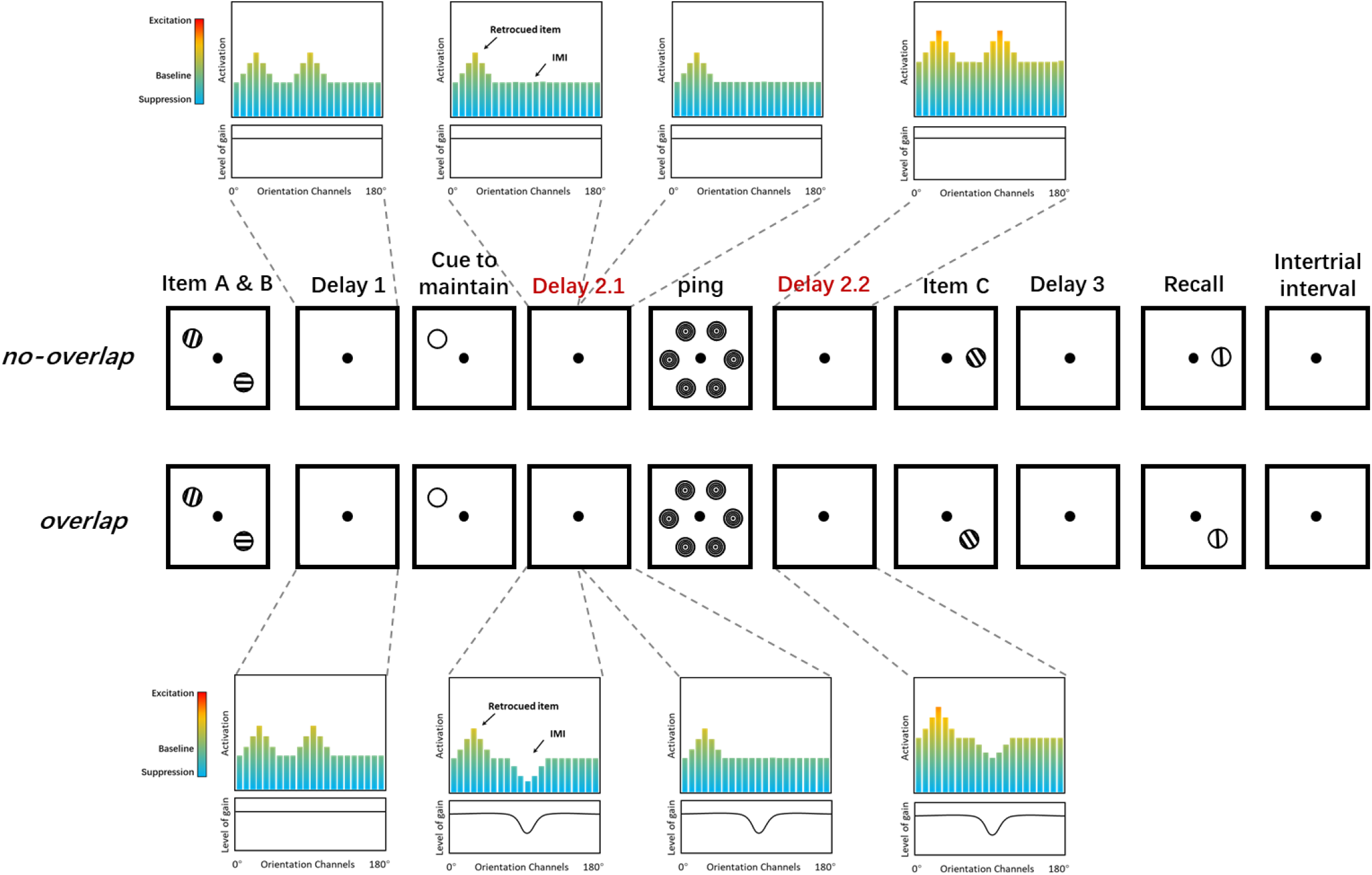
The hypothesized states of perceptual circuits that encode and maintain stimulus information in WM during different epochs of the trial. The panels above and below the timelines each have two elements: the colored bars represent the activation levels of hypothetical orientation channels and the black lines represent their level of gain. In the *no-overlap* trial, each sample item is represented during Delay 1 as elevated activity in the orientation channels corresponding to their value. During Delay 1.2, the representation of *A* remains elevated but that of *B* drops to baseline. Importantly, the activity-silent representation of *B* remains (not shown). Early in Delay 2.2, although the ping nonspecifically raises the activity level in every orientation channel, it also produces a reactivation of *B* (due to the persistence of *B*’s activity-silent representation). In the *overlap* trial, the cuing of *A* prompts a decrease in the gain of the channels corresponding to *B* plus a weak phasic activation of these channels, and the effect during Delay 1.2 is two-fold: a “suppressed” activity-based representation of *B* and the removal of the activity-silent representation of *B* (not shown). Later during Delay 2.1, the modified gain field persists but is not evident with only baseline levels of activity. Finally, early in Delay 2.2, responses to the ping reveal the modified gain field.

The pattern of results summarized in Figure 2 (formalized in the statement of *Preregistered hypotheses* in *Methods*) will provide neural evidence that the active removal of information from WM can be accomplished via a mechanism of adaptation-hijacking. It will also provide evidence relevant for accounts of the repulsive serial bias that is sometimes observed with perceptual discrimination tasks (e.g., Fritsche et al., 2020; Pascucci et al., 2019; Shan and Postle, Registered Report). Importantly, our design will also allow us to assess evidence for alternative accounts of active removal, and so deviations from the predicted pattern of results might nonetheless be informative. One possible alternative outcome is a failure to reconstruct the IMI during Delay 2.1 and Delay 2.2 (Figure 1) in the *overlap* condition. This would be consistent with the unbinding account of active removal, whereby the association between the content of the memory item and its context is actively broken (Lewis-Peacock et al., 2018). A different alternative pattern of results is that Delay 2.1 reconstructions of the IMI might have positive slopes in both conditions, but smaller in the *overlap* relative to the IMI in the *no-overlap* condition. This would suggest that there may not exist a mechanistic difference between what we have characterized as “active” versus “passive” removal from WM, and that the two only differ in terms of the magnitude of their effects. Finally, independent of the outcomes of stimulus-related analyses, we will assess evidence for the WMEM model’s prediction of larger mental context shifts on *overlap* versus *no-overlap* trials by assessing pattern similarity between Delay 1 and the late portion of Delay 2.1, in early visual cortex and in the medial temporal lobe (MTL; Figure 1). In the *overlap* condition, WMEM predicts a larger shift of mental context such that this discrepancy between mental contexts can compensate for the elevated level of cue competition.

## Methods

### Preregistered hypotheses

We propose to test 7 primary hypotheses in this Registered Report:

#### Hypothesis 1a

In the *overlap* condition, the reconstruction of the orientation of the IMI at TR 7, with an IEM trained on the retrocued item at TR 7, will have a significantly negative slope.

#### Hypothesis 1b

In the *no-overlap* condition, the reconstruction of the orientation of the IMI at TR 7, with an IEM trained on the retrocued item at TR 7, will be unsuccessful (i.e., slope not different from 0).

#### Hypothesis 1c

The slopes from 1a and 1b will differ.

#### Hypothesis 2a

In the *overlap* condition, if an IEM can be successfully trained to reconstruct the retrocued item at TR 12, the reconstruction of the orientation of the IMI at TR 12, with that same IEM, will be unsuccessful (i.e., slope not different from 0).

#### Hypothesis 2a’ (if needed)

In the *overlap* condition, if an IEM cannot be successfully trained to reconstruct the retrocued item at TR 12, the reconstruction of the orientation of the IMI at TR 12 with an IEM trained on the retrocued item at TR 7 will be unsuccessful (i.e., slope not different from 0).

#### Hypothesis 2b

In the *no-overlap* condition, if an IEM can be successfully trained to reconstruct the retrocued item at TR 12, the reconstruction of the orientation of the IMI at TR 12, with that same IEM will be unsuccessful (i.e., slope not different from 0).

#### Hypothesis 2b’ (if needed)

In the *no-overlap* condition, if an IEM cannot be successfully trained to reconstruct the retrocued item at TR 12, the reconstruction of the orientation of the IMI at TR 12 with an IEM trained on the retrocued item at TR 7 will be unsuccessful (i.e., slope not different from 0).

#### Hypothesis 3a

In the *overlap* condition, if an IEM can be successfully trained to reconstruct the retrocued item at TR 15+16, the reconstruction of the orientation of the IMI at TRs 15 + 16, with that same IEM, will have a significantly negative slope. *Hypothesis 3a’ (if needed)*: In the *overlap* condition, if an IEM cannot be successfully trained to reconstruct the retrocued item at TR 15+16, the reconstruction of the orientation of the IMI at TRs 15 + 16 with an IEM trained on the retrocued item at TR 7 will have a significantly negative slope.

#### Hypothesis 3b

In the *no-overlap* condition, if an IEM can be successfully trained to reconstruct the retrocued item at TR 15+16, the reconstruction of the orientation of the IMI at TRs 15 + 16 with that same IEM will have a significantly positive slope.

#### Hypothesis 3b’ (if needed)

In the *no-overlap* condition, if an IEM cannot be successfully trained to reconstruct the retrocued item at TR 15+16, the reconstruction of the orientation of the IMI at TRs 15 + 16, with an IEM trained on the retrocued item at TR 7 will have a significantly positive slope.

#### Hypothesis 3c

The slopes from 3a and 3b will differ.

#### Hypothesis 3c’ (if needed)

The slopes from 3a’ and 3b’ will differ.

#### Hypothesis 4a

In the in the early visual ROI the correlation coefficient between the high-dimensional activity pattern atTR 4 and the activity pattern at TR 12 will be higher in the *no-overlap* condition than in the *overlap* condition.

#### Hypothesis 4b

In the in the MTL ROI the correlation coefficient between the high-dimensional activity pattern at TR 4 and the activity pattern at TR 12 will be higher in the *no-overlap* condition than in the *overlap* condition.

### Summary Table

Note: There are two research questions: Hijacked-activation (HA) model and working memory episodic memory (WMEM) model; the sampling plan is the same for all hypotheses (n = 30)

| Hypothesis<br>(research<br>question) | Analysis | Pre-specified outcomes |
| --- | --- | --- |
| 1a (HA) | Bootstrapping | significantly negative slope = confirmed; slope not different from 0<br>suggests the breaking of content-context association of the IMI (Lewis-Peacock et al., 2018) as the mechanism of active removal |
| 1b (HA) | Bootstrapping | slope not different from 0 = confirmed; slope different from 0 (either<br>direction) means procedure failed to elicit expected initial signature of<br>passive removal |
| 1c (HA) | Bootstrapping | 1a and 1b differ = consistent with qualitative difference between active<br>and passive removal; no statistical difference suggests no mechanistic<br>difference between “active” and “passive” (e.g., perhaps a quantitative<br>but not qualitative difference) |
| *2a (HA)* | Bootstrapping | Slope not different from 0 = evidence for an active trace of the IMI is absent, a necessary condition for interpreting the effect of the ping; slope different from 0 may complicate interpretation of the effect of the ping. |
| *2a' (HA)* | Bootstrapping | This is an alternative way to assess *2a*, in the event that an IEM can NOT be successfully trained to reconstruct the retrocued item at TR 12;<br>interpretation of outcomes is the same as *2a* |
| *2b (HA)* | Bootstrapping | Same as *2a*, except for *no-overlap* condition |
| *2b' (HA)* | Bootstrapping | Same as *2a'*, except for *no-overlap* condition |
| *3a (HA)* | Bootstrapping | Negative slope = evidence for a key prediction of HA, the adaptation-like gain modulation of orientation-tuned sensory channels; positive slope = disconfirmation of HA; not different from 0 = failure to find evidence for HA, but consistent with the breaking of content-context association of the IMI (Lewis-Peacock et al., 2018) |
| *3a' (HA)* | Bootstrapping | This is an alternative way to assess *3a*, in the event that an IEM can NOT be successfully trained to reconstruct the retrocued item at TRs 14+15;<br>interpretation of outcomes is the same as *3a* |
| *3b (HA)* | Bootstrapping | Positive slope = evidence for a residual activity-silent representation of the IMI, an assumed consequence of passive removal; not different from 0 = failure to replicate Bae & Luck (2019) and Barbosa, Stein et al. (2020) |
| *3b' (HA)* | Bootstrapping | This is an alternative way to assess 3b, in the event that an IEM can NOT be successfully trained to reconstruct the retrocued item at TRs 14+15; interpretation of outcomes is the same as 3b |
| *3c (HA)* | Bootstrapping | *3a* and *3b* differ = consistent with qualitative difference between active and passive removal; no statistical difference suggests no mechanistic difference between “active” and “passive” (e.g., perhaps a quantitative but not qualitative difference) |
| *3c' (HA)* | Bootstrapping | This is an alternative way to assess 3c, in the event that either *3a'* or *3b'* were needed |
| *4a (WMEM)* | Permutation test | Correlation coefficient greater for *no-overlap* than for *overlap* in early visual ROI = support WMEM model; no difference between conditions = no support for WMEM model in early visual ROI |
| *4b (WMEM)* | Permutation test | Correlation coefficient greater for *no-overlap* than for *overlap* in MTL ROI = support WMEM model; no difference between conditions = no support for WMEM model in MTL ROI |

### Subjects

30 subjects who are 18-35 years in age with normal or corrected-to-normal vision and report no history of neurological disease will be recruited from the University of Wisconsin–Madison community. Informed consent will be obtained. All experimental procedures for the Registered Report have been approved by the University of Wisconsin– Madison Health Sciences Institutional Review Board.

#### Power analysis

Using data from Yu, Teng, and Postle (2020), in which a negative slope of the IEM reconstruction of the UMI in a DSR-of-orientation task has been observed, power analysis of the 2-tailed one sample *t*-test shows we will need data from 30 subjects to achieve 90% power to detect a significantly negative slope for the reconstruction of orientation of the UMI (Cohen’s d = 0.617), and data from 26 subjects to detect a significantly positive slope for the reconstruction of orientation of the PMI (Cohen’s d =0.675).

To the best of our knowledge, there is no established way to perform power analysis for bootstrapping, which we will use in the current study to test for the predicted positive and negative slopes of reconstructions. We used data from Yu, Teng, and Postle (2020) to simulate the *p*-values obtained from *t*-tests versus from bootstrapping with different sample sizes. Because this sample had data from 13 subjects, we generated estimates ranging from *N* = 8 to *N* =12, by randomly drawing N subjects from the sample, without replacement, and conducting a *t*-test and a bootstrap analysis on these data. For the *t*-tests, we collapsed over channel responses on both sides of the target channel, averaged them, and calculated the slope of the averaged UMI reconstruction of each subject with linear regression. The slopes were then compared to 0 with a 2-tailed one sample *t*-test. For bootstrapping, the method was the same as specified in the *Statistical Analyses* subsection of fMRI Analyses section of the *Methods*. This process was repeated 10 times at each N to get 10 (different) sets of subjects and 10 *p*-values for each test. For *N*=13, one *p*-value was obtained from each test. Across sample sizes, the bootstrapping was generally more sensitive than the *t*-test (Figure 3). Based on this, we reason that the sample size estimated by the power analysis for a *t*-test provides a conservative estimation of the sample size required in the current study (because we will be using the more sensitive bootstrapping procedure). In the current study we will use a sample size of 30 subjects.

**Figure 3.**
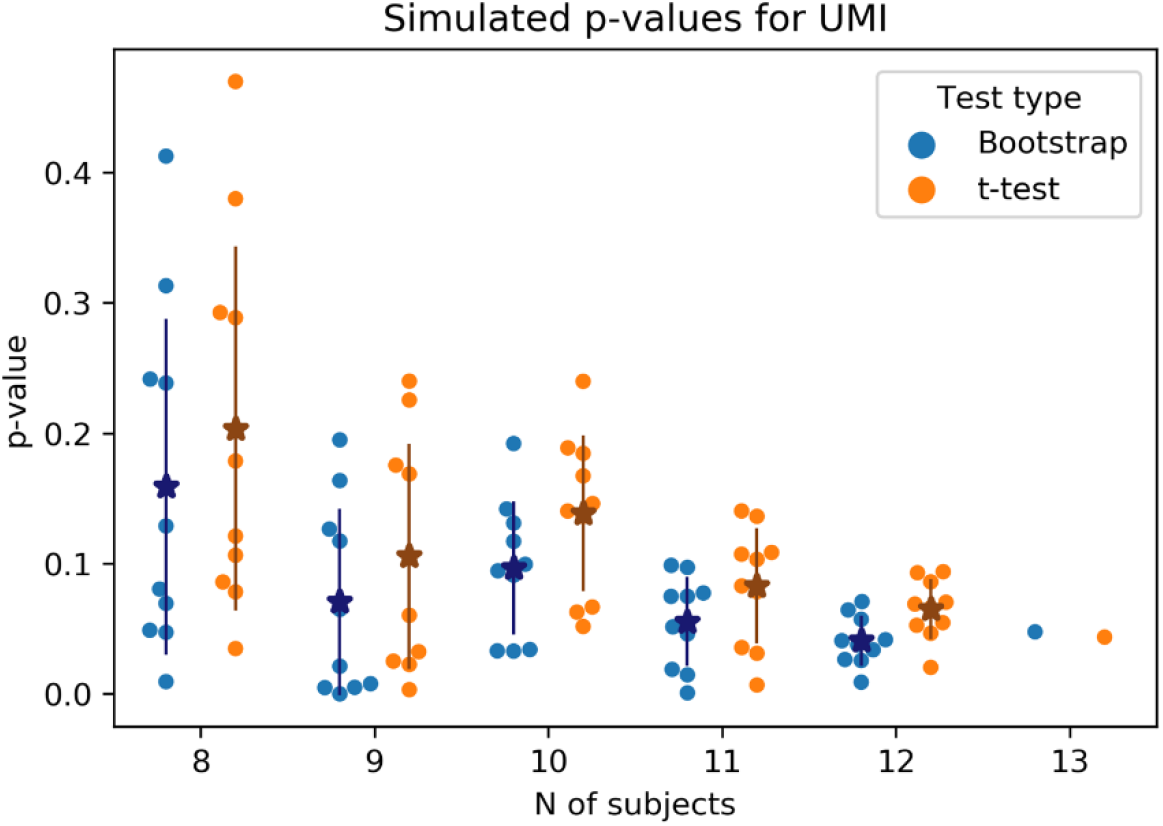
The *p*-values obtained from bootstrapping tests and *t*-tests of subsets of subjects from Yu, Teng, and Postle (2020). The darker stars overlaid on the dots represent the mean and the error bars show the standard deviation of each set of *p*-values.

### Stimuli and procedure

The stimulus presentation and response collection will be implemented with MATLAB (MathWorks, Natick, MA, USA) with the Psychtoolbox-3 extensions (Brainard, 1997; Pelli, 1997). The display will be projected into the scanner and onto a mirror mounted on the head coil at 60-Hz (Avotec Silent Vision 6011 projector; Avotec, Stuart, FL, USA). The viewing distance will be roughly 69 cm and the screen width will be 33.02 cm. The sample stimuli will be grayscale sinusoidal gratings (radius = 5°; contrast = 0.6; spatial frequency = 0.5 cycles/°; random phase angle) presented on gray background (L= 52, a = 0, and b = 0 in CIEL*ab space). There will be six possible sample orientations: 20°, 50°, 80°, 110°, 140°, 170°; with a random jitter of ±0°-3° added with each presentation. These and all ensuing stimuli will appear at any of six possible locations on an imaginary circle centered on fixation (radius of 8°, locations centered at each of these polar angle:s 30°, 90°, 150°, 210°, 270°, 330°). The retrocue will be a white circle (thickness=0.08°) with the same radius as the sample stimuli. Ping stimuli will be high contrast concentric circles with the same radius and spatial frequency as the gratings (contrast=1). The response dial will be a black bar (thickness=0.08°) corresponding to the diameter of a black circle with the same radius as the gratings (thickness=0.08°). Subjects will be instructed to adjust the orientation of the bar using an MR-compatible trackball (Current Designs, Philadelphia, PA, USA) and to report their response by pressing a button on the trackball when the orientation of the bar matches their memory for the probed sample. A white fixation dot will be present throughout each block (i.e., also during the ITI).

Each trial of ABC retrocuing will start with the simultaneous presentation of two samples (*A* and *B*; 1 s) followed by *Delay 1* (7 s). Next the retrocue will appear for 0.75 s at the location that had been occupied by either *A* or *B*, thereby designating a PMI (which might be tested at the end of the trial) and, by implication, the IMI (no longer relevant for that trial). The retrocue will be followed by *Delay 2*.*1* (15.25 s), which will be followed by the simultaneous presentation (0.25 s) of ping stimuli at each of the six locations, then *Delay 2*.*2* (7.75 s), then sample item *C* (1 s), then *Delay 3* (1 s). Finally, the response dial will appear at the location that had been occupied by the retrocued item or by item *C*, prompting the recall of the orientation of that item (4-s response window). The inter-trial interval ITI will vary randomly between 6, 8, and 10 s.

On each trial the orientation of items *A* and *B* will be randomly selected, with replacement, from the pool of six possible values. The locations of item *A* and *B* will be randomly selected from the six possible locations. To fully cross the orientations of item *A* and *B*, 21 unique trials are required. 252 trials (12 repetitions per unique trial) will be used for each condition. The retrocuing of *A* or *B* will be randomly determined on every trial. The orientation of item *C* will be randomly selected from the pool of six possible values (i.e., independent of *A* and *B*), and its location will depend on condition: in the *overlap* condition it will appear at the location that had been occupied by the uncued item; in the *no-overlap* condition it will appear in a location randomly selected from the four that had not been occupied by *A* or *B*. The retrocued item or the item *C* will be probed for recall equiprobably.

Trials will be blocked by condition (*overlap, no-overlap* condition), and subjects will be explicitly informed of the condition before the start of each block. Each subject will participate in 4 scanning sessions. The first scanning session will consist of 6 runs, each run corresponding to a 14-trial block. The three remaining scanning sessions will each consist of 10 runs (each run corresponding to a 14-trial block). There are fewer runs in the first session due to acquisition of structural images. To facilitate the consistent use of active removal and passive removal, within each session the first 3 blocks (for the first session) or 5 blocks (for the last three sessions) will be of one condition and the remaining blocks will be of the other condition. The order of conditions within a session will be counterbalanced across sessions and across subjects. In the first session, each subject will first do two practice blocks (one block for each condition) outside the scanner and another practice block (with the same condition as the first real block) inside the scanner. An Avotec RE-5700 eye-tracking system (Avotec) will be used to track eye position throughout each scanning session, and to assure that subjects’ eyes are open during the ping.

### Behavioral Data Analysis

The mean absolute error of recall across subjects will be calculated for each condition separately. The performance across the two conditions will be compared with a paired *t*-test.

### fMRI Data Acquisition

Whole-brain images will be acquired at the Lane Neuroimaging Laboratory at the University of Wisconsin–Madison HealthEmotions Research Institute (Department of Psychiatry) using a 3 Tesla GE MR scanner (Discovery MR750; GE Healthcare, Chicago, IL, USA). A high-resolution T1 image will be acquired with a fast spoiled gradient recalled echo sequence (8.2 ms TR, 3.2 ms TE, 12°flip angle, 176 axial slices, 256 × 256 in-plane, 1.0 mm isotropic) for each session. Functional data will be acquired with a gradient-echo echo-planar sequence (2 s repetition time [TR], 22 ms echo time [TE], 60°flip angle) within a 64 × 64 matrix (42 axial slices, 3 mm isotropic).

### fMRI Data Preprocessing

fMRI data will be preprocessed with the Analysis of Functional Neuroimages (AFNI) package (https://afni.nimh.nih.gov). To achieve a steady state of tissue magnetization, the first four TRs of each run will be discarded. The data will then be registered to the final volume of each scan and then to the anatomical images from the first session. Volumes will be motion corrected with six nuisance regressors to account for mead motion artifacts. Linear, quadratic, and cubic trends will be removed for each run and the z-scores of fMRI time series data will be calculated within each run.

### fMRI Analyses

#### Task-related activity

The fMRI data will be fitted to a general linear model (GLM) with regressors for each epoch of the task -- *Encoding A&B* (2 s), *Delay 1* (6 s), *Delay 2*.*1* (16 s), *Delay 2*.*2* (8 s), *Encoding C* (2 s), *Recall* (4 s) – each convolved with a canonical hemodynamic response function, as well as nuisance covariates for between-trial and between-scan drift, and head motion.

#### ROI creation

Hypothesis tests will be carried out in “early visual cortex” ROIs. First, an anatomical ROI of early visual cortex will be created from masks corresponding to V1 and V2 (merged, both hemispheres), taken from the probabilistic atlas of Wang and colleagues (2015) and warped to each subject’s structural scan in native space. Hypothesis testing will be carried out in the 500 voxels within the anatomical early visual cortex ROI with have the strongest weights on the Encoding A&B regressor, which we refer to as the early visual ROI. The MTL ROI will be manually selected for each subject.

#### Inverted Encoding modeling

IEM analyses will be performed with custom functions in MATLAB. In IEM, the responses of each voxel are assumed to be a weighted sum of responses of several hypothetical tuning channels. Six tuning channels of orientation will be used and the tuning curve of each channel will be defined as a half-wave–rectified sinusoid raised to the eighth power. We will first compute the weight matrix W (v voxels × k channels) that projects the hypothesized channel responses C1 (k channels × n trials) to the measured voxel responses B1 (v voxels × n trials) with the training dataset to get the estimate of the weight matrix *Ŵ*. Then we use *Ŵ* to reconstruct the channels responses 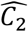 from the voxel activities B2 of the testing dataset. The relationship between B1, W and C1 will be characterized by

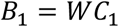

The least-squared estimate of the weight matrix (*Ŵ*) will be calculated using linear regression:

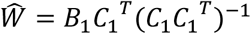

The channels responses 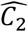of the testing dataset will then be calculated with the weight matrix (*Ŵ*) and the BOLD data (B2):

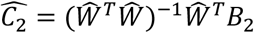

The IEMs will be trained with the orientation of the retrocued item and test on the orientation of the retrocued item or the IMI at each TR after the offset of item A and B and before the onset of item C. We will use a leave-one-run-out procedure in which the model will be trained with data of all but one run and tested on the left-out run. This process will be repeated until the reconstruction of all runs is acquired. The estimated channels responses will be centered on the orientation of the tested item. The reconstruction will be generated on all TRs but our pre-registered hypotheses will focus on specific TRs: *Hypothesis 1a, 1b and 1c* will focus on TR 7; *Hypothesis 2a and 2b* will focus on TR 12. For *Hypothesis 3a, 3b and 3c*, the averaged BOLD of TR 15 and TR 16 will be used to train and test the model. In case we cannot get a reliable reconstruction of the retrocued item at TR 12 and/or TRs 15+16 due to the representation of the retrocued item shifts to an activity-silent state after a long time span since the presentation of the item, the retrocued item at TR 7 will be used to train the model and TR 12 (for *Hypothesis 2a and 2b*) and/or TRs 15+16 (for *Hypothesis 3a, 3b and 3c*) will be tested.

#### Statistical Analyses

The strength of IEM reconstructions of memory items will be operationalized by their slope. We will collapse over channel responses on both sides of the target channel, average them, and calculate the slope of the reconstruction with linear regression for each subject separately.

We will use bootstrapping to test the statistical significance of the slope of each reconstruction (Ester et al., 2016, 2015). We will randomly sample 30 reconstructions from the pool of averaged reconstructions of each subject, with replacement, and calculate the average of the channel responses. This process will be repeated 10,000 times to get 10,000 average reconstructions, and the slopes of these reconstructions will be calculated. Two-tailed *p*-values will be computed as the proportion of positive or negative slopes, whichever is smaller, multiplied by 2.

To test the difference between slopes, we will calculate the difference between the 2 slopes of interest for each one of the 10,000 resampled data sets. Two-tailed *p*-values will be the proportion of positive or negative differences, whichever is smaller, multiplied by 2.

#### Shift of mental context

Within each trial, we will calculate the Pearson correlation coefficient between the activity in the early visual ROI at TR 4 and the activity in the early visual ROI at TR 12. An average correlation coefficient will be calculated for each subject in each condition. The correlation coefficients will then be averaged across subjects in each condition and we’ll calculate the difference of these two coefficients (*r*_no-overlap_ -*r*_overlap_). We will use a one-tailed permutation test to test whether the correlation coefficient is significantly higher in the no-overlap condition than in the overlap condition. We will randomly exchange the condition label of each subject’s average coefficient, calculate the average r across subjects for each condition, and calculate the difference of the coefficients. This process will be repeated 50,000 times to get 50,000 differences under the null hypothesis that there is no difference in correlation between the *overlap* condition and the *no-overlap* condition. One-tailed *p*-values will be computed as the proportion of differences that are higher than the real difference.

### Timeline

We anticipate the data collection will take about 1 year. We will carry out data processing and analysis in parallel to data collection as new data are collected. The analyses of data and write-up and submission of the Stage 2 report are expected to be completed within 4 months after all data are collected. If there are pandemic-related interruptions, the project will be delayed accordingly.

